# Converting Muscle-mimetic Biomaterials to Cartilage-like Materials

**DOI:** 10.1101/2021.05.18.444710

**Authors:** Linglan Fu, Lan Li, Bin Xue, Jing Jin, Yi Cao, Qing Jiang, Hongbin Li

## Abstract

Load-bearing tissues, such as muscle and cartilage, exhibit mechanical properties that often combine high elasticity, high toughness and fast recovery, despite their different stiffness (∼100 kPa for muscles and one to several MPa for cartilage).^1-7^ The advance in protein engineering and protein mechanics has made it possible to engineer protein-based biomaterials to mimic soft load-bearing tissues, such as muscles.^8-10^ However, it is challenging to engineer protein biomaterials to achieve the mechanical properties exhibited by stiff tissues, such as articular cartilage,^6,11^ or to develop stiff synthetic extracellular matrices for cartilage stem/progenitor cell differentiation^12^. By employing physical entanglements^13^ of protein chains and force-induced protein unfolding,^14,15^ here we report the engineering of a highly tough and stiff protein hydrogel to mimic articular cartilage. By crosslinking an engineered artificial elastomeric protein from its unfolded state, we introduced chain entanglement into the hydrogel network. Upon renaturation, the entangled protein chain network and forced protein unfolding entailed this single network protein hydrogel with superb mechanical properties in both tensile and compression tests, showing a Young’s modulus of ∼0.7 MPa and toughness of 250 kJ/m^3^ in tensile testing; and ∼1.7 MPa in compressive modulus and toughness of 3.2 MJ/m^3^. The energy dissipation in both tensile and compression tests is reversible and the hydrogel can recovery its mechanical properties rapidly. Moreover, this hydrogel can withstand a compression stress of >60 MPa without failure, amongst the highest compressive strength achieved by a hydrogel. These properties are comparable to those of articular cartilage, making this protein hydrogel a novel cartilage-mimetic biomaterial. Our study opened up a new potential avenue towards engineering protein hydrogel-based substitute for articular cartilage, and may also help develop protein biomaterials with superb mechanical properties for applications in soft actuators and robotics.

---

Many load-bearing tissues, ranging from muscle to cartilage, are protein-based biomaterials that exhibit finely regulated mechanical properties to uniquely suit their biological functions.^1,16^ To engineer biomimetics of these biological tissues, protein-based hydrogels have been widely explored.^17^ Protein hydrogels are generally soft, with a Young’s modulus up to ∼100 kilopascal (kPa).^8,18^ Thus, current protein hydrogel technologies have achieved considerable success in mimicking softer tissues^17-19^, such as muscle.^8,9,15^ However, stiffer tissues often have a modulus on the order of megapascal (MPa) and bear tensile as well as compressive loads, making them challenging to achieve for the current protein hydrogel technology. For example, articular cartilage is a superb load-bearing material made of collagen and proteoglycans. It has a modulus ranging from one to several MPa, can withstand a load up to a hundred MPa and sustain millions of cycles of loading-unloading without much fatigue, showing fast recovery in its shape and mechanical properties.^1,7,20^ Engineering protein hydrogels that can combine these seemingly mutually incompatible properties (high stiffness, high toughness and fast recovery) remains challenging.

In order to engineer protein hydrogels with such a combination of mechanical features, higher crosslinking density and efficient microscopic mechanism for energy dissipation and recovery are essential. Articular cartilage realized this unique combination of mechanical features by using a protein fiber-proteoglycan composite approach.^21^ However, this approach is difficult to design from bottom up.^22,23^ Conversely, muscles may provide an inspiration for an alternative approach for engineering cartilage-like protein-based materials. Forced-unfolding and refolding of globular proteins is a mechanism employed in muscle to allow for effective energy dissipation upon overstretching, and recovery upon relaxation.^3,14,24^ This mechanism has been used to engineer protein hydrogels to mimic the passive elastic properties of muscles.^8-10,15^ We reason that if an efficient method can be developed to significantly improve the stiffness of muscle-mimetic biomaterials, one should be able to take advantage of the forced-unfolding as an efficient energy dissipation mechanism to convert muscle-mimetic biomaterials into biomaterials with mechanical properties resembling those of articular cartilage.

Chain entanglement is an important strengthening mechanism in polymeric materials.^13^ Due to their long length, polymer chains in the network can cross each other, resulting in chain entanglement. Chain entanglement effectively increases the crosslinking density in the polymer network and leads to improved Young’s modulus.^13^ However, in muscle fibers, the muscle protein titin are organized as parallel bundles without chain entanglement.^2,3^ In the engineered muscle mimetic protein hydrogels constructed from tandem modular protein-based elastomeric proteins, no chain entanglement is present either, due to the short contour length of such elastomeric proteins (about 10 to 40 nm in length).^8,15^ Here we explore the feasibility to impart the soft, muscle-mimetic protein biomaterial with chain entanglement to significantly increase its stiffness. For this purpose, we used the elastomeric protein (FL)_8_ as a model system.

FL is a de novo designed ferredoxin-like globular protein.^25^ Single molecule optical tweezers experiments showed that FL is mechanically labile, and unfolds and refolds readily at ∼5 pN (Fig. S1).^15^ The elastomeric protein (FL)_8_ was used to engineer highly stretchy and tough protein hydrogels, in which the forced-unfolding of FL domains served as a highly effective means in dissipating energy in the hydrogel.^15^ However, the Young’s modulus of the (FL)_8_ hydrogel is only ∼15 kPa.^15^ We sought to use (FL)_8_ as a model system for enhancing its mechanical stiffness.

The molecular weight of (FL)_8_ is ∼80 kDa, but its contour length in its native state is only ∼10 nm. Thus, there is no chain entanglement in the native (FL)_8_ hydrogels. However, in its unfolded state, (FL)_8_ is ∼260 nm long, showing the characteristic length of a polymer chain. In the concentrated solution of unfolded (FL)_8_ (>150 mg/mL), the unfolded polypeptide chains will overlap and likely entangle.^26^ We reason that if unfolded (FL)_8_ is crosslinked from its concentrated solution, inter-chain entanglement could be trapped by the chemical crosslinks in the crosslinked hydrogel network.

To test this feasibility, the well-developed [Ru(bpy)_3_]^2+^-mediated photochemical crosslinking strategy, which crosslinks two tyrosine residues in proximity into a dityrosine adduct, was employed to construct (FL)_8_ hydrogels.^8,18,27,28^ We used the DC (<u>d</u>enatured <u>c</u>rosslinking) method to construct the denatured (FL)_8_ hydrogels, in which the unfolded (FL)_8_ was photochemically crosslinked into hydrogels directly from its concentrated solution (20%, 200 mg/mL) in 7 M guanidine hydrochloride (GdHCl). The as-prepared hydrogel was then equilibrated in 7 M GdHCl to obtain the <u>d</u>enatured DC hydrogel (referred to as the D-DC hydrogel). As a control, we also constructed a denatured (FL)_8_ hydrogel that is free of chain entanglement using the NC (<u>n</u>ative <u>c</u>rosslinking) method (Fig. S2). In this method, native (FL)_8_ solution (200 mg/mL) in phosphate buffered saline (PBS) was first crosslinked into a hydrogel, which was free of chain entanglement due to the short length of native (FL)_8_. Then the prepared hydrogel was denatured and equilibrated in 7M GdHCl to obtain the <u>d</u>enatured NC hydrogel (referred to as the D-NC hydrogel).

Fig. 1A shows the photographs of both D-DC and D-NC (FL)_8_ hydrogels prepared using the same ring-shaped mold as well as their stress-strain curves. Evidently, the D-DC hydrogel was self-standing and swelled to a much less degree than the D-NC hydrogel, while the D-NC hydrogel ring-shaped sample collapsed onto itself. The D-DC hydrogel displayed a Young’s modulus of 56 kPa, significantly higher than that of D-NC hydrogel (∼1 kPa).

**Figure 1.**
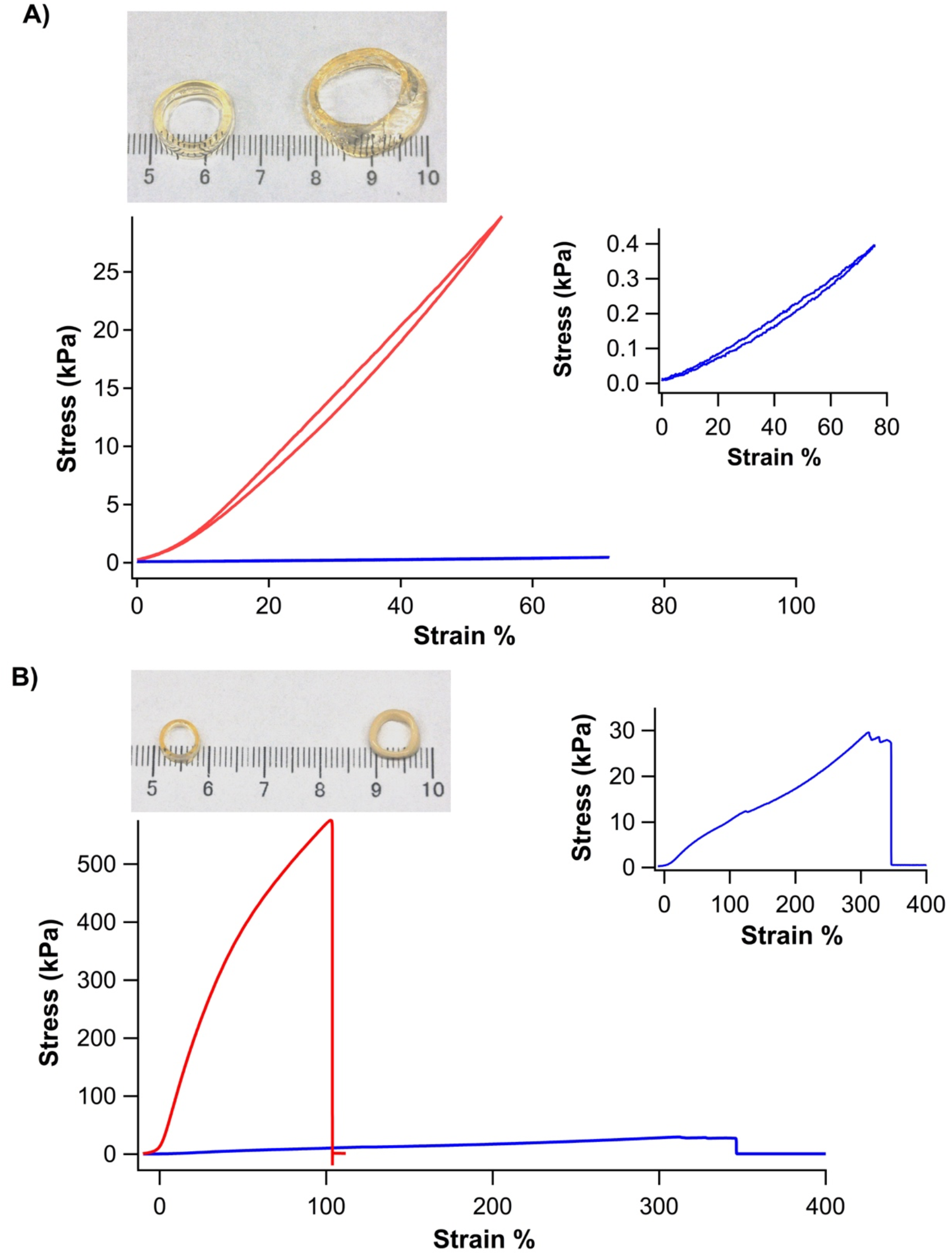
Physical entanglements enhanced the stiffness of the (FL)_8_ hydrogels. A) Stress-strain curves of D-DC (red) and D-NC (blue) (FL)_8_ hydrogels (200 mg/mL). Inset is the zoom view of the stress-strain curve of the D-NC hydrogel. The D-DC hydrogel showed a Young’s modulus of ∼50 kPa, significantly higher than that of D-NC hydrogel (∼1 kPa). Top panel shows the photographs of both hydrogels after being equilibrated in 7M GdHCl. The D-DC hydrogel is self-standing and swells to a much less degree than the D-NC hydrogel. B) Stress-strain curves of N-DC (red) and N-NC (blue) (FL)_8_ hydrogels (200 mg/mL). Inset is the zoom view of the stress-strain curve of the N-NC hydrogel. The D-DC hydrogel showed a Young’s modulus of ∼0.7 MPa, significantly higher than that of N-NC hydrogel (∼20 kPa). The N-DC hydrogel ruptured at ∼100% strain. Top panel shows the photographs of both hydrogels equilibrated in PBS. The N-DC hydrogel is translucent, while N-NC hydrogel is opaque.

According to the classical rubber elasticity theory,^13^ *G=ρRT/M*_*c*_, where *G* is the shear modulus, *ρ* is the density of the polymer in the network, *Mc* is the average molecular weight between two neighboring crosslinking points, *R* is the gas constant and *T* is temperature, the stiffness (Young’s modulus) of a hydrogel network is determined by the crosslinking density. Thus, our results suggested a much higher effective crosslinking density for the D-DC gel than the D-NC gel. To examine if these two preparation methods led to different number of dityrosine crosslinking points in the (FL)_8_ hydrogel, we quantified the number of dityrosine adducts by measuring the characteristic dityrosine fluorescence (Fig. S3A).^18,29^ Our results showed that both hydrogels contained roughly the same number of dityrosine crosslinking points (∼17% of the total number of tyrosine residues in FL domains were cross-linked into dityrosine adducts). Since the D-DC and D-NC hydrogels had the same number of crosslinking points, the same unfolded protein chain and the same protein chain-solvent interaction, the higher effective crosslinking density of the D-DC hydrogel should originate from additional effective crosslinking points resulted from the chain entanglements of unfolded (FL)_8_ polypeptide chains in the D-DC hydrogel network. Evidently, the chain entanglement significantly enhanced the stiffness of the denatured (FL)_8_ hydrogels.

Chain entanglement in a chemically crosslinked hydrogel network cannot be removed without disrupting the chemical crosslinking network. Thus, chain entanglement will be retained in the hydrogel network even if the polypeptide chains undergo conformational changes, such as folding. Taking advantage of this unique topological feature, we “ renatured” the D-DC hydrogel in PBS buffer to allow the refolding of FL domain to obtain the <u>n</u>ative DC (N-DC) (FL)_8_ hydrogel.

We observed that after renaturing in PBS, the N-DC hydrogel deswelled dramatically compared with the D-DC hydrogel, and changed from being transparent to largely translucent (Fig. 1B). In contrast, upon renaturation, the N-NC hydrogel deswelled and became opaque. The swelling ratio of the N-DC hydrogel was smaller than that of the N-NC hydrogel. Moreover, both hydrogels can be cycled between their native and denatured states (N-DC to D-DC, N-NC to D-NC) for many cycles without noticeable change in their respective physical appearances and properties (Fig. S4). The deswelling is likely due to the refolding of some FL domains in PBS, which is accompanied by a significant shortening of the contour length of the polyprotein (from 260 nm to 10 nm). Microscopically, both N-DC and N-NC hydrogels showed microporous structures, but the mesh size of N-DC hydrogel was significantly smaller than that of the N-NC hydrogel, consistent with the smaller swelling ratio of the N-DC hydrogel (Fig. S5).

During renaturation, due to the presence of chain entanglement and the shortening of the polypeptide chain, it is expected that some FL domains would not be able to fold into their native state. Instead, they likely assumed a hydrophobically collapsed state and/or random coil state, as PBS is a poor solvent for the unfolded FL domain. Therefore, it is likely that the N-DC (FL)_8_ hydrogel network assumed a single network structure consisting of folded FL domains as well as unfolded ones, as schematically shown in Fig. 2.

**Figure 2.**
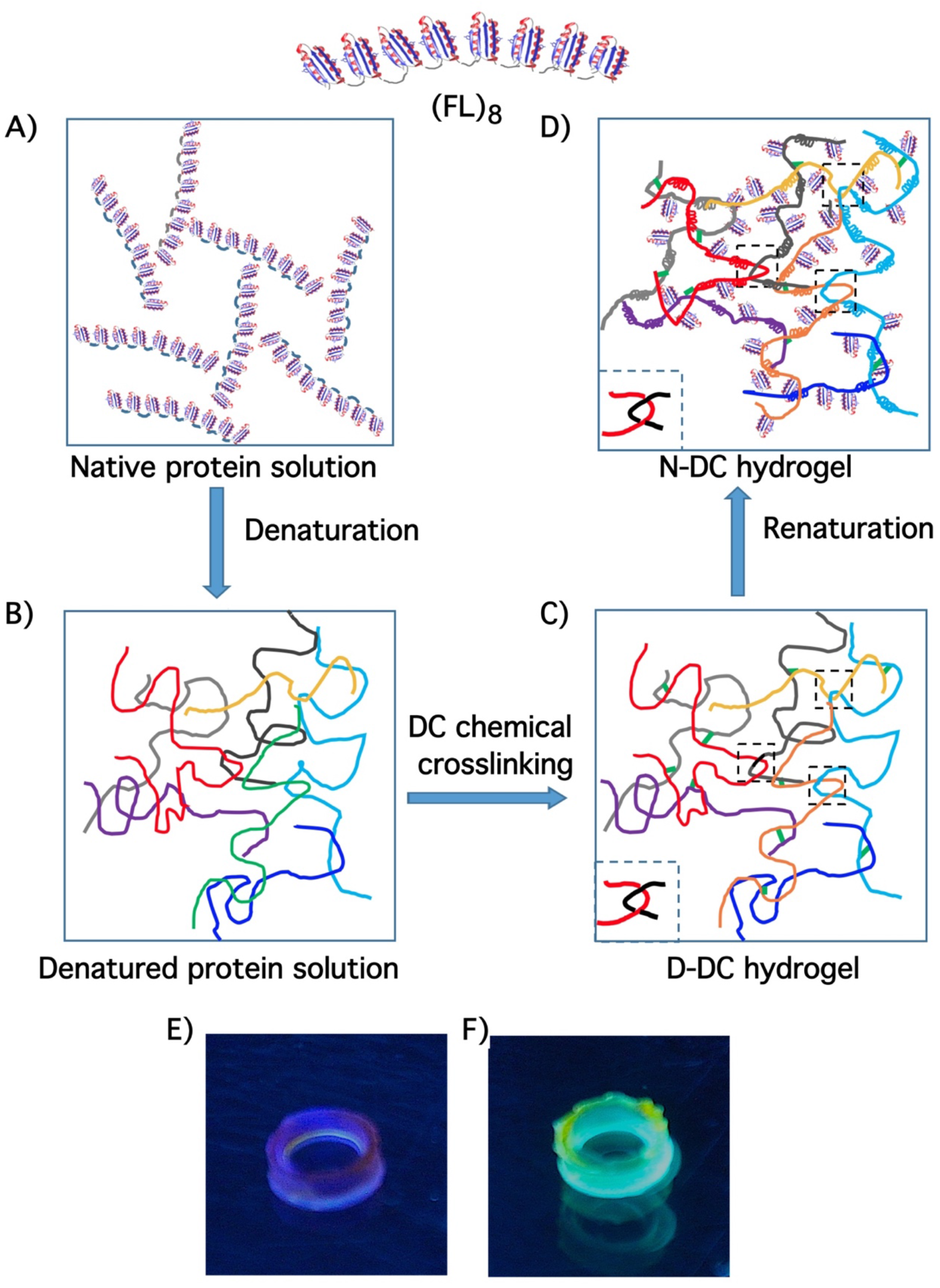
Schematics of the chain entangled network structure of the N-DC (FL)_8_ hydrogel. A-D) Schematics of the preparation of D-DC and N-DC (FL)_8_ hydrogels. The elastomeric protein (FL)_8_ was first dissolved in PBS to high concentration (∼200 mg/mL) (A). When denatured in GdHCl, the unfolded (FL)_8_ polypeptide chain behaved as random coils, which overlapped with one another due to high protein concentration, leading to possible chain entanglements (B). Upon photochemical crosslinking, chain entanglements were retained or created, leading to a network of entangled polypeptide chains, in the D-DC hydrogel (C). Entangled chains are highlighted in dashed squares. Lower left bottom shows a zoomed view of one such chain entanglement. When the D-DC hydrogel was equilibrated in PBS, some FL domains refolded, while others underwent hydrophobic collapse. In the renatured hydrogel (N-DC hydrogel), the chain entanglements remained, which are highlighted in dashed squares (D). For clarity, different molecules in B-D) are colored differently. E-F) Photographs of (FL-M23C)_8_ N-DC hydrogels. E) a (FL-M23C)_8_ N-DC hydrogel under UV illumination light. The blue fluorescence was from the dityrosine crosslinking points; F) a (FL-M23C)_8_ N-DC hydrogel under UV-illumination after labeling with IAEDANS. The cyan fluorescence was from the labeling of the exposed cysteine residues, and indicated that some FL domains were unfolded in the hydrogel.

To verify the existence of unfolded FL domains in the N-DC hydrogel, we used a well-established cysteine shotgun labeling approach, which allows for labeling of only solvent-exposed cysteine residues using the thiol reactive fluorescent dye 5-((2-((iodoacetyl)amino)ethyl)amino)naphthalene-1-sulfonic acid (IAEDANS).^30^ For this, we used a cysteine variant FL-M23C, where the buried residue Met23 in FL was mutated to Cys. Cys23 is sequestered in the hydrophobic core of the folded FL and can only be labeled with IAEDANS when FL-M23C is unfolded.^29^ The (FLM23C)_8_-based hydrogels showed similar physical and mechanical properties as (FL)_8_ (Fig. S6). After IAEDANS labeling in PBS, the N-DC hydrogel showed the characteristic cyan fluorescence of IAEDANS under UV illumination (Fig. 2E-2F and Fig. S3B). Quantitative analysis showed that ∼50% of the FL domains were unfolded in the N-DC hydrogel. In contrast, only ∼23% of the FL domains were unfolded in the N-NC hydrogel.^29^ These results indicated that the N-DC hydrogel is indeed a single network hydrogel consisting of folded and unfolded FL domains.

We then carried out tensile and compression tests to measure the mechanical properties of the N-DC (FL)_8_ hydrogel. A typical stress-strain curve of the N-DC hydrogel in PBS buffer is shown in Fig. 1B, where the N-DC hydrogel was stretched to failure. For comparison, the stress-strain curve of the N-NC hydrogel is also shown. It is evident that the stiffness of the N-DC hydrogel increased dramatically compared with that of the N-NC hydrogel and D-DC hydrogel. The N-DC hydrogel displayed a Young’s modulus of 0.70 ± 0.11 MPa (n=18) (Fig. 1B and Fig. S7A), significantly higher than that of protein-based hydrogels reported in the literature.^8,9,18,19,31^ The breaking strain of the N-DC hydrogel is 107 ± 14%, indicative of its good stretchability (Fig. S7A). Similar Young’s modulus and breaking strain were also observed for 15% N-DC (FL)_8_ hydrogel as well as N-DC hydrogels constructed from (FL)_16_. However, 10% N-DC (FL)_8_ hydrogel showed similar Young’s modulus but much smaller breaking strain (50%) (Fig. S8).

Moreover, during cyclic stretching-relaxation experiments, the N-DC hydrogel exhibited a large hysteresis, indicative of a significant amount of energy dissipation at high strains during stretching (Fig. 3A). The average energy dissipation of the N-DC hydrogel is 250 ± 68 kJ/m^3^, demonstrating its superb toughness (Fig. S7B). Moreover, the energy dissipation of the N-DC (FL)_8_ hydrogel was fully reversible, and the hysteresis can be recovered rapidly once the hydrogel was relaxed to zero strain (Fig. 3B). After being stretched to 60% strain and then relaxed to zero strain, 70% of the hysteresis was recovered right after the hydrogel was relaxed, and the rest 30% hysteresis recovered more slowly following a double-exponential kinetics, with a rate constant k_1_ of 0.05 ± 0.02 s^-1^ and k_2_ of (1.7 ± 0.3)×10^−3^ s^-1^, respectively (Fig. 3C). This reversible hysteresis is likely due to the forced-unfolding, and subsequent refolding of the FL domains in the hydrogel network, consistent with the fast refolding rate of FL domain at zero force.^15^

**Figure 3.**
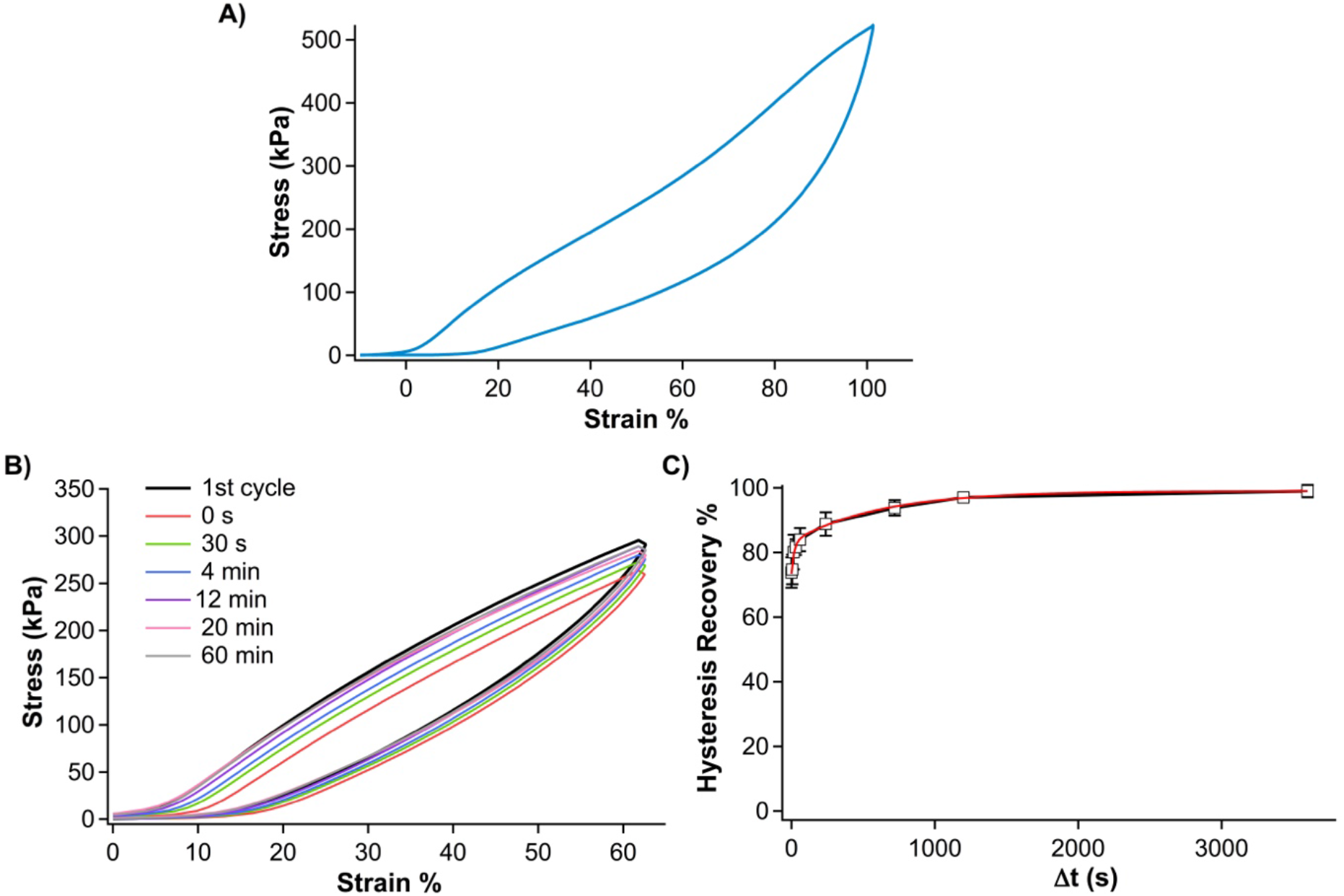
The N-DC (FL)_8_ hydrogels can dissipate a large amount of energy. A) Stretching-relaxation stress-strain curves of the N-DC (FL)_8_ hydrogel. A large hysteresis was present in the stretching and relaxation curves, indicative of large energy dissipation. B) The hysteresis between stretching and relaxation curves can be recovered rapidly. The hydrogel was first stretched to ∼60% strain and then relaxed to zero strain. After waiting for certain time Δt, the hydrogel was subject to the stretching-relaxation cycle again. The hysteresis recovery can be directly observed. C) The kinetics of the hysteresis recovery in N-DC (FL)_8_ hydrogel. ∼70% of the hysteresis can be recovered rapidly within a few seconds, and the rest 30% hysteresis can be recovered following a double-exponential kinetics. The red line is a double exponential fit to the data, with a rate constant k_1_ of 0.05 ± 0.02 s^-1^ and k_2_ of (1.7 ± 0.3)×10^−3^ s^-1^, respectively.

Evidently, the N-DC (FL)_8_ hydrogels displayed tensile mechanical properties that uniquely combined a high Young’s modulus (∼0.7 MPa), high toughness as well as fast recovery, a combination that is difficult to achieve as individual properties are often mutually incompatible. Additionally, the high Young’s modulus and toughness of this single network protein hydrogel are amongst the highest of the engineered protein hydrogels, and even compare favorably with those of some specially designed synthetic polymer hydrogels of special network structures, such as double network hydrogels,^32,33^ or polymer composite hydrogels.^22^

In addition to these superb tensile properties, the N-DC (FL)_8_ hydrogel demonstrated even more striking compressive properties. We found that the N-DC (FL)_8_ hydrogel is super tough and can resist slicing with a sharp scalpel, despite that it contains ∼60% water (Fig. 4A). To quantify the compressive mechanical properties of the N-DC (FL)_8_ hydrogel, standard compression tests were carried out (Fig. 4B). The stress-strain curves showed that the N-DC (FL)_8_ hydrogels displayed a compressive modulus of ∼1.7 MPa at 10-20% strain. In comparison, the N-NC (FL)_8_ hydrogels only showed a compressive modulus of ∼50 kPa (Fig. 4B, inset), again revealing the significant enhancement effect of chain entanglement on the stiffness of the (FL)_8_ hydrogels. As the strain increased, the stress of the N-DC (FL)_8_ hydrogel showed a much more pronounced increase: the modulus at 75% strain reached ∼10 MPa. Strikingly, the N-DC FL hydrogel could be compressed to more than 80% strain and sustain a compressive stress of as high as 75 MPa without fracture (Fig. S9). The average compressive strength of the N-DC (FL)_8_ hydrogel is 68 ± 12 MPa (n=7), and an average failure strain of 82 ± 3%. At the failure strain, a small crack often started to appear in the hydrogel sample. However, the failure was not brittle, as evidenced by the subsequent stress-strain curves of the same hydrogel sample (Fig. S10). These results demonstrated the super high compressive strength of the N-DC (FL)_8_ hydrogel. The compressive strength of the N-DC (FL)_8_ hydrogel is amongst the highest strength achieved by hydrogels (Table S1), and compares favorably with that of articular cartilage.^4,7,21^ As a comparison, the super tough double network polymer hydrogels typically fractured at a stress of no more than 20 MPa.^32,33^

**Figure 4.**
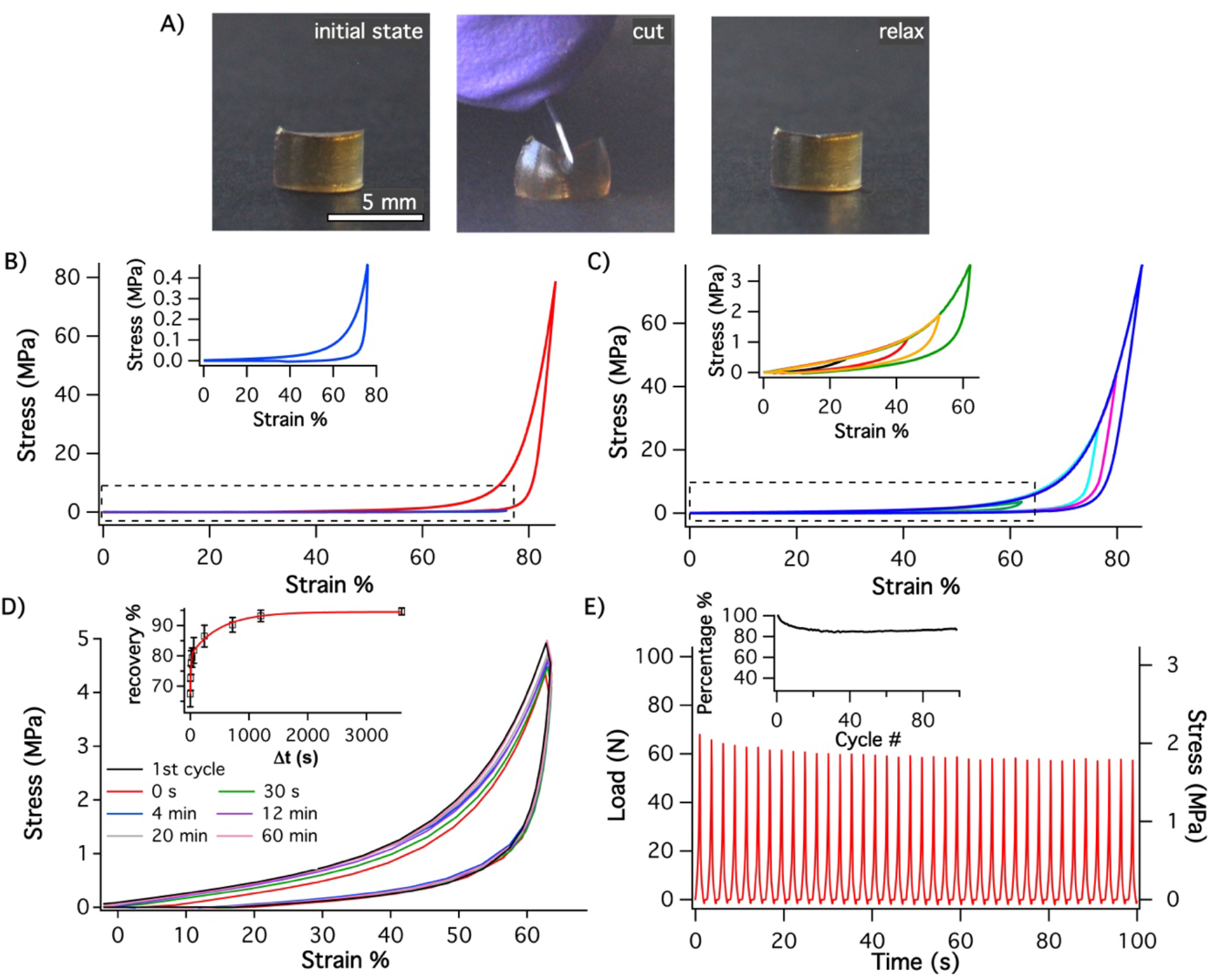
N-DC (FL)_8_ hydrogels displayed superb compressive mechanical properties. A) Photographs show that the N-DC (FL)_8_ hydrogel can resist cutting with a sharp scalpel. B) Compressive stress-strain curves of the N-DC (red) and N-NC (FL)_8_ (blue) hydrogels. Inset is a zoom view of the stress-strain curves of N-NC hydrogel. The N-DC hydrogel can be compressed to more than 80% strain and sustain a compressive stress of >70 MPa without failure. A large hysteresis was present between the loading and unloading curves, indicating that a large amount of energy was dissipated. C) Consecutive compression-unloading curve of the N-DC hydrogel. The hysteresis grows with the increasing of the strain. The toughness of the hydrogel is ∼3.2 MJ/m^3^. The inset is a zoom view of the stress-strain curves at lower strain. D) Consecutive compression-unloading cycles show that the hysteresis of the N-DC hydrogel can be recovered rapidly. Inset shows the hysteresis recovery kinetics of the hydrogel. ∼65% hysteresis can be recovered right after unloading, and the rest hysteresis can be recovered following a double exponential kinetics, with k_1_ of 0.10 ± 0.02 s^-1^ and k_2_ of (2.0 ± 0.3)×10^−3^ s^-1^. E) Consecutive loading-unloading curves of the N-DC (FL)_8_ hydrogel at a frequency of ∼0.37 Hz. The pulling speed was 100 mm/min. In each cycle, the hydrogel was stretched to 60% strain and subsequently relaxed to zero strain. After 100 cycles, the hydrogel displayed little fatigue, and the stress of the hydrogel at 60% strain retained ∼90% of the original stress in the first cycle.

In the compression-unloading cycles, a large hysteresis was observed (Fig. 4C), indicative of a large amount of energy being dissipated during compression. The hysteresis increased as the strain increased. At 80% strain, the hydrogel exhibited a toughness of 3.2 ± 0.6 MJ/m^3^ (n=7). Moreover, the (FL)_8_ hydrogel displayed superb recovery properties (Fig. 4D and Movie S1). At lower strains (<40%), the hydrogel recovered its dimension right at the end of the loading-unloading cycles, suggesting that the recovery of the hydrogel is fast. Even at a large compression strain (>60%), ∼65% of the hysteresis can be recovered right after unloading. The remaining 35% hysteresis could be recovered within an hour following a double exponential kinetics (Fig. 4D). Furthermore, after 100 consecutive loading-unloading cycles at a frequency of 0. 37 Hz and a final strain of 50%, the consecutive loading-unloading curves did not show much change, and the stress of the N-DC (FL)_8_ hydrogel at 50% strain retained ∼90% of its original stress (Fig. 4E). Similar results were obtained at a frequency of 0.67 Hz and 0.08 Hz (Fig. S11), demonstrating that the N-DC (FL)_8_ hydrogel can quickly recover its mechanical properties and do not show much fatigue.

Collectively, these results revealed that the N-DC (FL)_8_ hydrogels are mechanically strong and tough, and can recover their shape and mechanical properties rapidly and do not show much mechanical fatigue. Moreover, these protein hydrogels showed excellent long term stability: after being stored in PBS (with 0.2‰ NaN_3_) for over six months, their physical shape and mechanical properties remained largely unchanged. These exceptional mechanical properties and their unique integration in one material are rare for protein hydrogels, and compare favorably with those of polymer hydrogels with special network structure (Table S1). These properties closely reproduced many features of articular cartilage, and thus make the N-DC (FL)_8_ hydrogels a cartilage-mimetic protein biomaterial.

These superb mechanical properties of N-DC (FL)_8_ hydrogels are likely resulted from a combination of factors, including chain entanglement, folded and hydrophobically collapsed FL domains in the hydrogel network, as well as the forced unfolding and refolding of FL domains. These unique features likely make this DC method for protein hydrogelation broadly applicable. Indeed, as shown in Table S2, in protein hydrogels constructed from a range of elastomeric proteins, which range from all α proteins to α/β proteins, we significantly enhanced their stiffness via this DC hydrogelation method and improved their Young’s modulus to several hundred kPa, depending on the inherent tyrosine content of the elastomeric protein. Similar enhancement was also achieved in the compressive modulus of these protein hydrogels (Table S2). These results have demonstrated the generality of this novel method.

Stiff biological tissues, such as cartilage, tendons and ligaments, often integrate seemingly mutually incompatible mechanical properties into themselves.^1^ Mimicking such properties using synthetic hydrogels has been challenging. It often occurs that optimizing one property is at the expenses of another one. To alleviate this issue and achieve high stiffness and high toughness, polymer hydrogels of designed network structures and polymer composite hydrogels have been developed,^32-39^ such as double network hydrogel,^32,33^ co-joined network hydrogel^34^ and slide-ring hydrogel.^35,36^ Sacrificial bonds/weak secondary network that can be ruptured are often introduced into the hydrogel as an energy dissipation mechanism.^33,40,41^ Although high stiffness and high toughness have been achieved in some of these hydrogels, slow recovery and mechanical fatigue are often present, due to the irreversible rupture of these sacrificial bonds and/or slow dynamics of weak secondary networks. Although proteins are attractive building blocks to construct soft biomaterials, protein hydrogels are generally soft, making them largely inept to mimic stiff tissues (Table S1). Here we demonstrated a novel DC hydrogelation approach, which combines forced-unfolding of globular proteins and chain entanglement, to enable the engineering of strong and tough protein hydrogels. On the one hand, forced-unfolding of globular proteins provides an efficient mechanism for energy dissipation, and the ability to refold allows the hydrogel to recovery its mechanical properties rapidly and minimize mechanical fatigue. On the other hand, chain entanglement allows the hydrogel to achieve high stiffness. In so doing, these effects work cooperatively to allow the integration of high stiffness, high toughness, fast recovery and high compressive strength into protein hydrogels. Although the engineered protein hydrogel has a single network structure, its superb mechanical properties essentially convert a muscle-like soft biomaterials to a stiff material exhibiting mechanical properties that mimic cartilage. Our study has significantly expanded the range of mechanical properties that protein hydrogels can achieve, and thus make many existing protein hydrogels as potential candidates for developing cartilage-mimetic protein hydrogels. Given the generality of this novel approach and the richness of potential protein building blocks, our study may open up an exciting new area for exploration, as well as developing novel protein-based biomaterials for applications in fields ranging from cartilage repair to soft robotics and actuators.

## Methods Summary

The (FL)_8_ and (FL-M23C)_8_ polyproteins were engineered using previously published protocols.^29^ Protein hydrogels were constructed using a photochemical crosslinking strategy as described before. For the DC hydrogel, the photochemical crosslinking was carried out in 7 M GdHCl solution. For the NC hydrogel, the photochemical crosslinking was carried out in PBS solution. Tensile and compression tests were performed on an Instron-5500R universal testing system. The local slope at 15% strain on the loading curve was taken as the modulus for both tensile and compression tests. The toughness was calculated as the area between the loading curves at a given strain using custom-written software in IgorPro.

## Acknowledgement

This work is supported by the Natural Sciences and Engineering Research Council of Canada. We thank Prof. R. Shadwick for assistance with Instron tests.

## Supplementary Information

### Material and methods

#### Protein engineering

FL domain is a redesigned variant of Di-I_5 (PDB code: 2KL8).^1,2^ The amino acid sequence of FL is: MGEFDIRFRT DDDEQFEKVL KEMNRRARKD AGTVTYTRDG NDFEIRITGI SEQNRKELAK EVERLAKEQN ITVTYTERGS LE. The genes of the polyprotein ferredoxin-like proteins (FL)_8_, (FLM23C)_8_, (FL)_16_ were constructed following standard and well-established molecular biology methods as reported previouly.^3^ Other polyproteins, (GB1)_8_, (NuG2)_8_, GRG_5_RG_4_R, N_4_RN_4_RNR and (GA)_8_, were constructed following the same method. Polyprotein genes were inserted into the vector pQE80L for protein expression in *E. coli* strain DH5α. Seeding culture was allowed to grow overnight in 10 mL 2.5% Luria-Bertani broth (LB) medium containing 100 mg/L ampicillin. The overnight culture was used to inoculate 1 L of LB medium which was grown at 37 °C and 225 rpm for 3 hours to reach an OD_600_ of ∼0.8. Protein expression was induced with 1 mM isopropyl-1-β-D-thiogalactoside (IPTG) and continued at 37°C for 4 hours. The cells were harvested by centrifugation at 4000 rpm for 10 mins at 4 °C and then frozen at -80 °C. For polyprotein purification, cells were thawed and resuspended in 1x phosphate-buffered saline (PBS) and lysed by incubation with 1 mg/mL lysozyme for 30 mins. Nucleic acids were removed by adding 0.1 mg/mL of both DNase and RNase. The supernatant with soluble protein was collected after centrifuging the cell mixture at 12000 rpm for 60 mins. The soluble His_6_-tagged protein was purified using a Co^2+^ affinity column. The yields of (FL)_8_, (FLM23C)_8_, and (FL)_16_ are approximately 80 mg, 80 mg and 45 mg respectively per liter of bacterial culture. Purified proteins were dialyzed extensively against deionized water for 2 days to remove residual NaCl, imidazole, and phosphate. Then the protein solution was filtered and lyophilized, and stored at room temperature until use. Bovine serum albumin (BSA) lyophilized powder was purchase from Sigma-Aldrich.

#### Hydrogel preparation and dimension measurements

Lyophilized native (FL)_8_ protein was processed using two different gelation methods. The NC (<u>n</u>ative <u>c</u>rosslinking) hydrogels were prepared in native state by dissolving and gelating proteins in 1x PBS. After gelation, N-NC hydrogels were soaked in PBS, while the D-NC hydrogels were stored in 1xPBS containing 7M guanidine-hydrochloride (GdHCl) and achieved swelling equilibrium. The DC (<u>d</u>enatured <u>c</u>rosslinking) hydrogels (D-DC and N-DC) were prepared by dissolving the lyophilized (FL)_8_ in 7M GdHCl for 2 hrs before use. The denatured protein solution was crosslinked into hydrogels and equilibrated in 7M GdHCl to obtain D-DC hydrogels, while N-DC hydrogels were renatured in PBS on a rocker by changing fresh PBS ten times over the course of 1 day until reaching equilibrium.

Gelation of (FL)_8_, (FLM23C)_8_ and (FL)_16_ were based on photochemical crosslinking strategy described previously.^2-5^ To prepare 20 % (w/v) hydrogels, lyophilized proteins (200 mg/mL) were re-dissolved in PBS (D-NC and N-NC) or 7M GdHCl in PBS (D-DC and N-DC) respectively. A typical crosslinking reaction mixture contained 200 mg/mL of polyprotein, 50 mM ammonium persulfate (APS) and 200 μM [Ru(bpy)_3_]Cl_2_. The protein mixture was quickly cast into a custom-made plexiglass ring-shaped mold (d_in_=8 mm, d_out_=10 mm, h=3 mm), and was exposed to a 200 W fiber optical white light source placed 10 cm above the mold for 10 min at room temperature. After gelation was complete, the hydrogel sample was carefully taken out of the mold. After the ring-shaped hydrogels were stored in the desired buffers for 3 hrs (N-DC gels for 24 hrs), the outer diameter, thickness, width and weight of all ring-shape equilibrated swollen/deswelling samples were measured before tensile tests. For compressive tests and SEM imaging, the hydrogels were prepared in a cylindrical shape following the same gelation procedures. The hydrogels preparation and the tensile (E) and compressive (Y) moduli measurements of (GB1)_8_, (NuG2)_8_, GRG_5_RG_4_R, NRN_4_RN_4_R, (GA)_8_ and BSA followed the same procedures.

#### Swelling ratio and water content measurements

For swelling ratio and water content measurements, ring-shaped hydrogels were weighed immediately after being taken out of the mold, and the weight was recorded as *m*_*i*_. The swollen weight *m*_*s*_ was recorded after gently removing excess buffer from equilibrated hydrogels. To measure dry weight of the gels (*m*_*dry*_), the hydrogels were firstly immersed in deionized water to remove extra salts, then lyophilized for 2 days and dried in a 70 °C incubator for 1 day. The swelling ratio (*r*) of the hydrogels was calculated using the following formula,

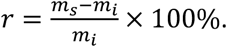

The water content (*w*) was determined by the dried gel (*m*_*dry*_) and the equilibrated gel (*m*_*s*_) (measured specimens, n=4)

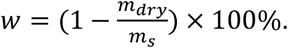

#### Tensile tests

Tensile tests were performed using an Instron-5500R tensometer with a custom-made force gauge and 5-N load transducer. The ring-shaped hydrogel specimen was stretched and relaxed in PBS (N-DC and N-NC) or 7 M GdHCl in PBS (D-DC and D-NC) at constant temperature (25 °C) without special preconditioning. The stress was calculated by dividing the load by the initial cross-sectional area of the hydrogel sample. The Young’s modulus, breaking strain, and energy dissipation were measured using an extension rate of 25 mm/min. The stress at 15% strain is taken as the Young’s modulus of the sample. Toughness was determined by integrating stress-strain curves where specimens were loaded directly to failure. Energy dissipation was calculated by integrating loop area between stretching and relaxing stress-strain curves. In hysteresis recovery experiments, a pulling rate of 200 mm/min was used. The same ring sample was stretched and relaxed with various time intervals.

#### Compression tests

Uniaxial compression tests were performed on cylinder-shaped hydrogels that were swollen to equilibrium using the Instron-5500R with 5000-N load transducer in air at room temperature. The dimensions (height (*h*_*0*_) and diameter (*d*_*0*_)) of the equilibrated N-DC and N-NC (FL)_8_ hydrogel samples are: *h*_*0*_: 3.0 mm and *d*_*0*_: 6.5 mm for the N-DC hydrogel, and *h*_*0*_: 5.0 mm and *d*_*0*_: 5.0 mm for the N-NC hydrogel. The gel was put on the lower plate, while the upper plate approached the sample slowly until a rise in force was detected, indicating contact between the platen and the gel. The stress was calculated by dividing the load by the initial cross-sectional area of the hydrogel sample. Unless a different rate is stated, the gel was compressed and relaxed at a compression speed of 2 mm/min. No water was squeezed out of the gels during compression. The compressive modulus was measured at a strain of 10-20 %. The maximum stress and strain were determined at failure points, where the first crack in the gel was observed. Energy dissipation was calculated by integrating loop area between compressing and relaxing stress-strain curves (n=7). In hysteresis recovery experiments, a compression rate of 100 mm/min was used.

#### Scanning electron microscopy (SEM) imaging

20 % (w/v) D-NC and N-NC (FL)_8_ hydrogel samples were prepared for SEM imaging using a Hitachi S4700 scanning electron microscope. The samples were then shock-frozen in liquid nitrogen, and quickly transferred to a freeze drier where they were lyophilized for 24 hrs. Lyophilized samples were then carefully fractured in liquid nitrogen, and fixed on aluminum stubs. The sample surface was coated by 8 nm of gold prior to SEM measurements.

#### Characterization of dityrosine cross-links in hydrogels by acid hydrolysis-fluorescence method

The degree of dityrosine crosslinking in both NC and DC (FL)_8_ hydrogels was characterized following a well-established fluorimetry method.^6^ Dityrosine emits at a wavelength of 410 nm when excited at 315 nm. For quantification of the dityrosine and dityrosine-like compounds generated in NC and DC (FL)_8_ hydrogels, we followed a well-established fluorescence standard curve method.^6^ Typically, 20 % (w/v) hydrogel samples (∼25 mg) were reacted with 100 μL HCl (6 N) in a sealed 1.5 mL centrifuge tube in a metal heat block at 105 °C for 2 hrs to achieve full hydrolysis of the peptide bonds. Then, 100 μL of acid hydrolysis product was transferred into a new 1.5 mL centrifuge tube and neutralized by 10 μL NaOH (5 M). Next, 100 mM Na_2_CO_3_-NaHCO_3_ buffer (pH 9.9) was added to a final volume of 1 mL. Fluorescence spectra of the samples were measured by a Varian Cary Eclipse fluorescence spectrophotometer. According to the fluorescence-concentration standard calibration curve of dityrosine, the yield of dityrosine and dityrosine-like products in the hydrogel was then determined (n=8).

#### Cysteine shotgun fluorescence labeling by IAEDANS and fluorescence measurements

DC and NC (FLM23C)_8_ hydrogels for cysteine shotgun labeling were prepared with the same protein concentration and gel preparation procedures as the wild-type (FL)_8_. The labeling reaction was performed in the dark at room temperature for 3 hrs in PBS buffer (pH 7.4) containing 5 mM TCEP and 2 mM 5-((2-[(iodoacetyl)amino]ethyl)amino)naphthalene-1-sulfonic acid (IAEDANS). As a control, D-DC and D-NC (FLM23C)_8_ hydrogels incubated in 7 M GdHCl (containing 5 mM TCEP, pH 7.4) were also labeled using IAEDANS. Then all hydrogels were transferred to PBS buffer containing 5 mM β-mercaptoethanol to quench the reaction. To remove excess labeling dye, additional PBS buffer (containing 5 mM β-mercaptoethanol) was added, and changed five times over the course of 5 hrs until fluorescence could no longer be detected in the buffer solution.

To quantify the fluorescence intensity of IAEDANS labeled hydrogels, we digested the hydrogels with trypsin at 37 °C for 5 hrs. The digestion reaction contained 5 % trypsin (relative to the hydrogel weight), 25 mM NH_4_HCO_3_, 10 mM CaCl_2_, 1 M GdHCl and 10 mM dithiothreitol. Unlabeled (FLM23C)_8_ hydrogel was digested in the same way to serve as a negative control. After digestion, 50 μL of the digested mixture was diluted to 3 mL using PBS buffer. The fluorescence emission spectrum was measured by a Varian Cary Eclipse fluorescence spectrometer using an excitation wavelength of 336 nm and emission at 490 nm was monitored. An IAEDANS standard calibration curve was also created, covering linear concentration range of 0-60 μM (n=8).

#### Single-molecule optical tweezers measurements

Single-molecule optical tweezers measurements were carried out using a MiniTweezers setup as previously.^7,8^ Sample preparation including the protein–DNA construct formation and force measurement protocols was adapted from protocols described previously.^8^ Force-distance curves of the protein–DNA construct were obtained using constant velocity pulling protocol.

---

**Figure S1.**
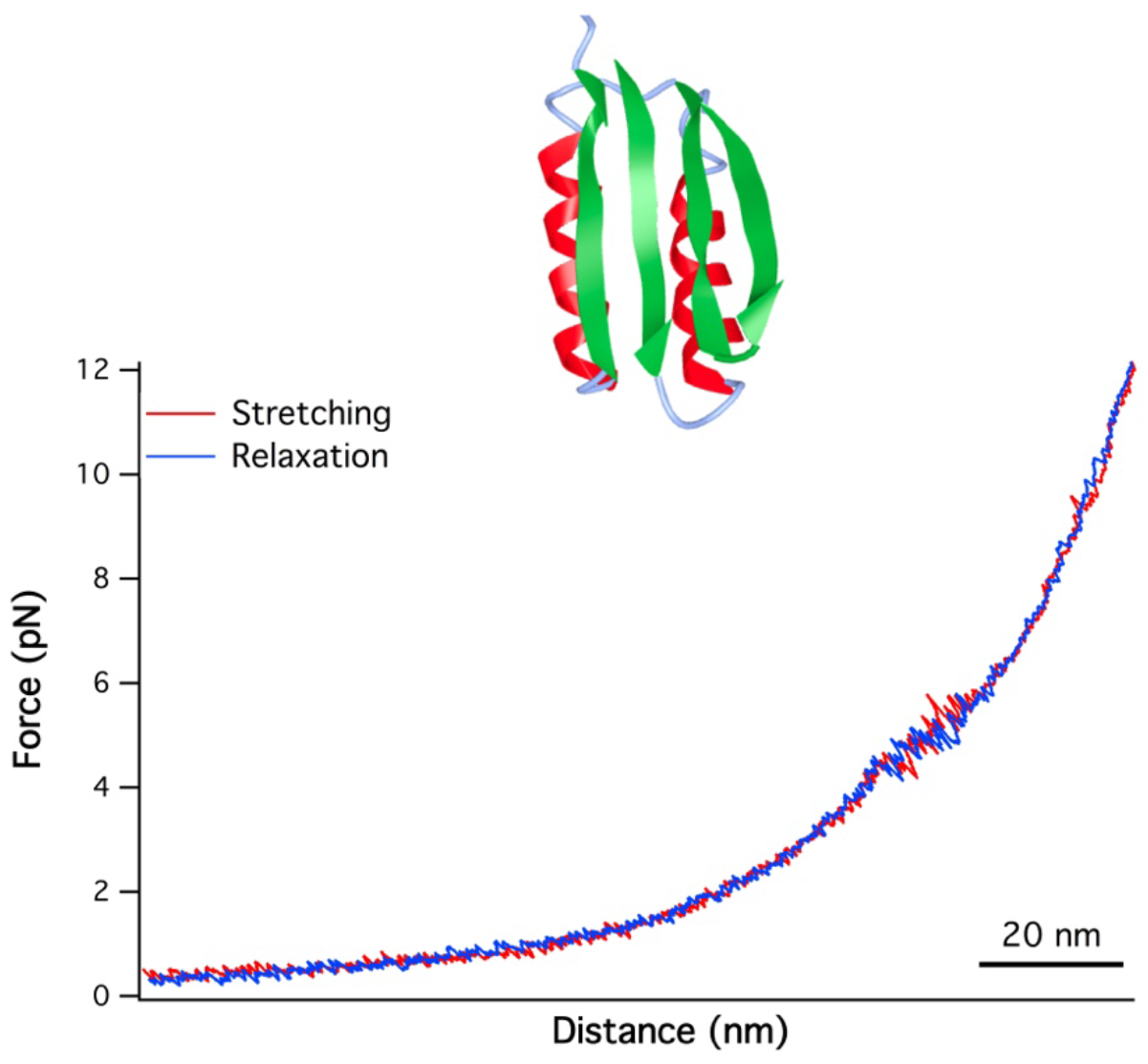
Representative force-distance curves of FL domain at a pulling speed of 50 nm/s. The unfolding-refolding of FL occurred at ∼ 5 pN, making FL a mechanically labile protein. The inset shows the three-dimensional structure of FL (PDB code: 2KL8).

**Figure S2.**
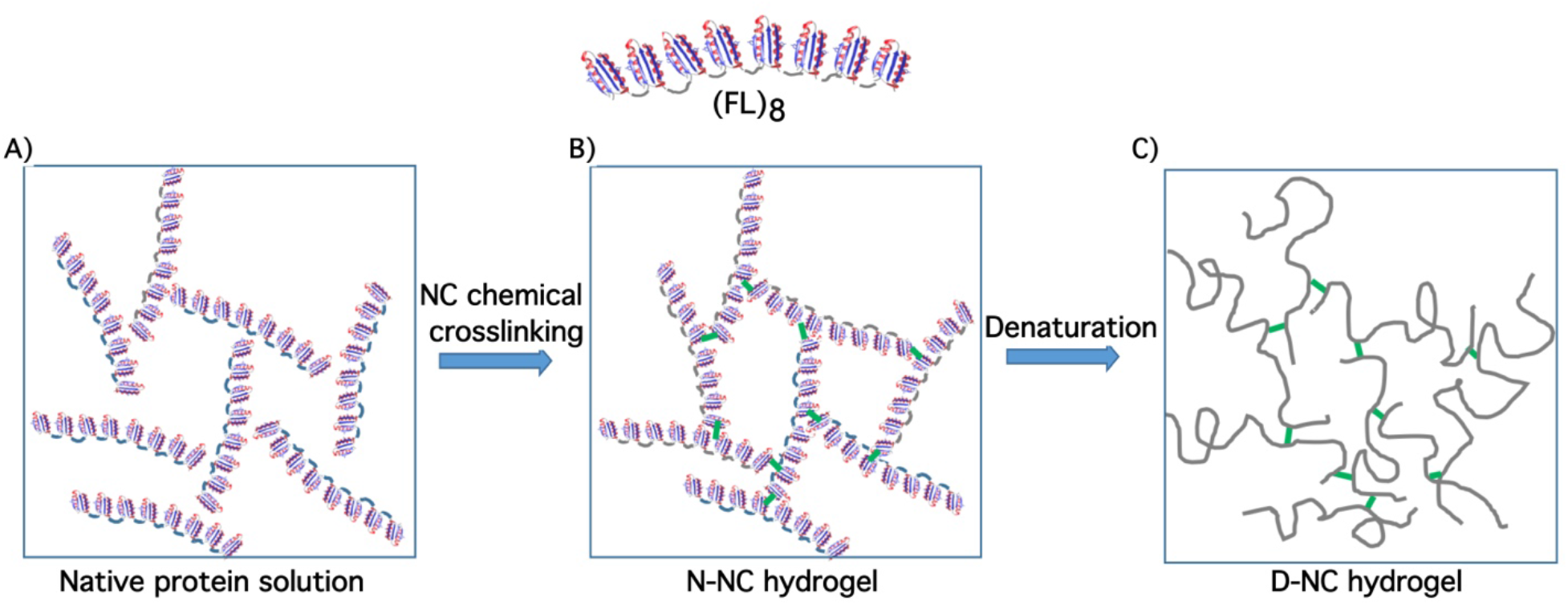
Schematics of the NC-(FL)_8_ hydrogels. The elastomeric protein (FL)_8_ was first dissolved in PBS to a high concentration (∼200 mg/mL) (A). Upon photochemical crosslinking, (FL)_8_ were crosslinked into a hydrogel network without chain entanglements, due to the short length of folded (FL)_8_, resulting in the N-NC hydrogel (B). When denatured in GdHCl, the (FL)_8_ in the hydrogel network unfolded, and behaved as random coils. The resultant D-NC hydrogel is also free of chain entanglement (C).

**Figure S3.**
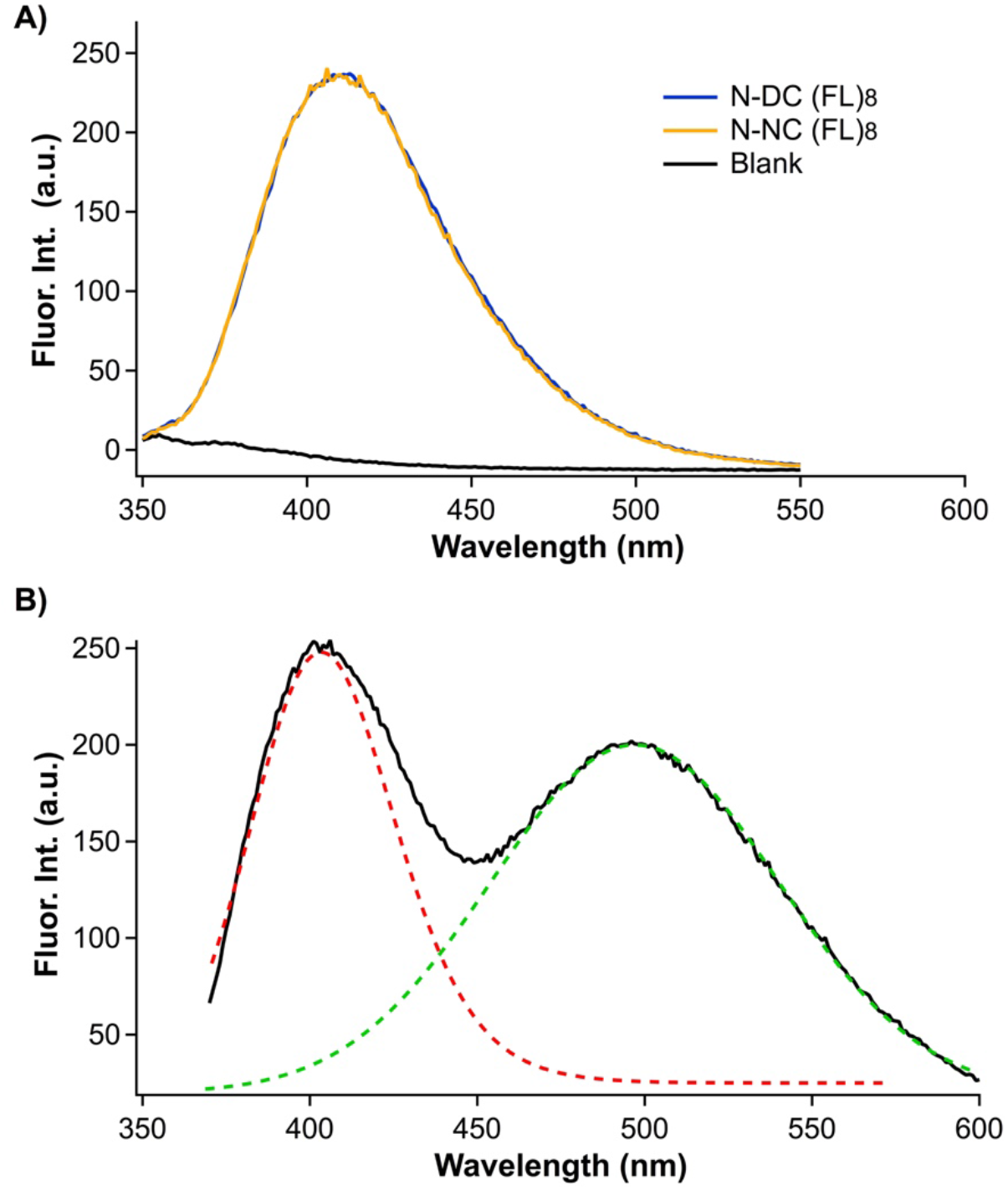
Fluorescence spectra of (FL)_8_ hydrogels. A) Fluorescence spectra of acid hydrolyzed 20% N-DC and N-NC (FL)_8_ hydrogels prepared from the same weight of lyophilized (FL)_8_ proteins. Fluorescence at 410 nm is resulted from the dityrosine fluorescence. B) Fluorescence spectrum of IAEDANS labeled 20% (FL-M23C)_8_ hydrogel. Dotted lines are Gaussian fits to the two fluorescence peaks, one is the dityrosine fluorescence at 410 nm, and the other one is the IAEDANS fluorescence at 490 nm.

**Figure S4.**
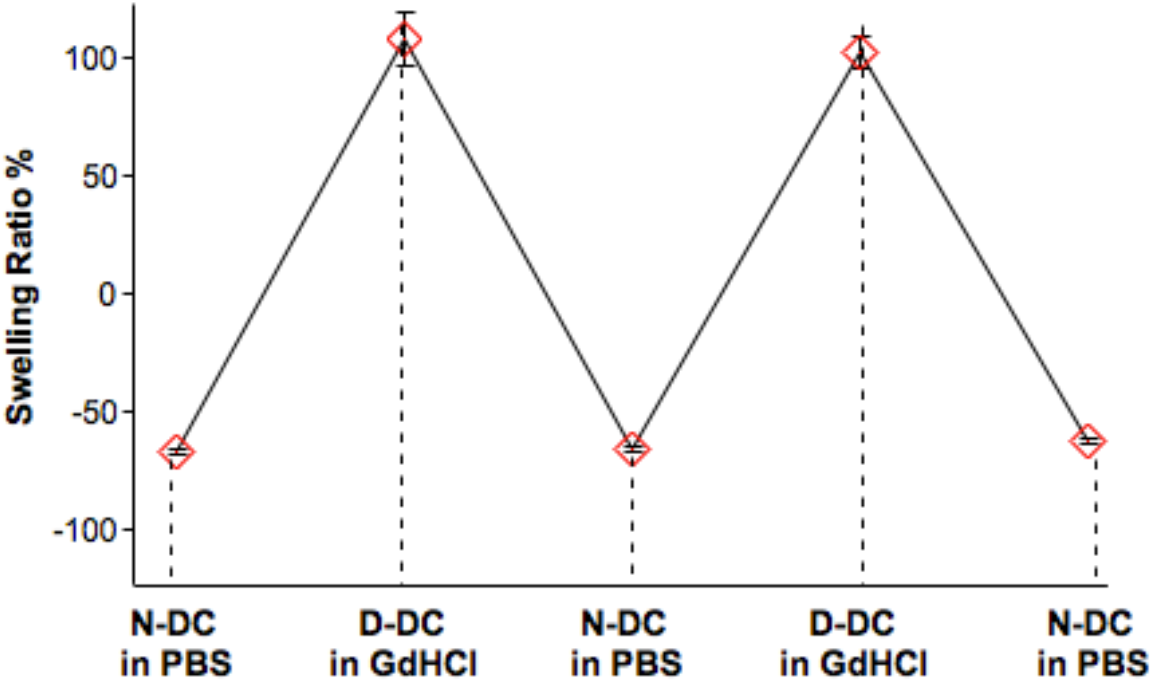
DC (FL)_8_ hydrogels can be cycled between its N-DC and D-DC states reversibly.

**Figure S5.**
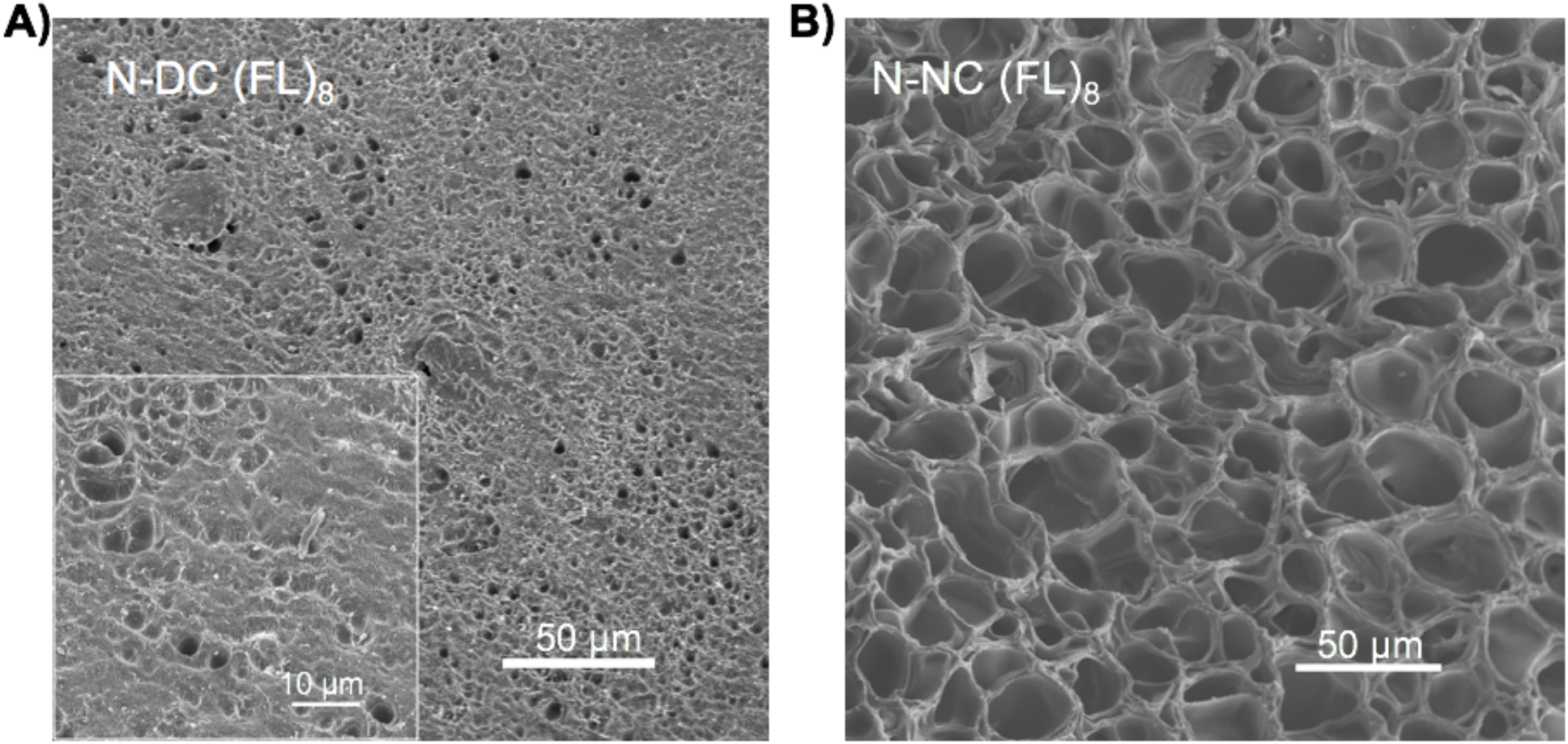
Photographs of scanning electron microscopy imaging of the N-DC (A) and N-NC (B) (FL)_8_ hydrogels. Both hydrogels showed porous network structures, however, the pore size of the N-DC hydrogel (∼2 μm) is significantly smaller than that of the N-NC hydrogel (∼20 μm).

**Figure S6.**
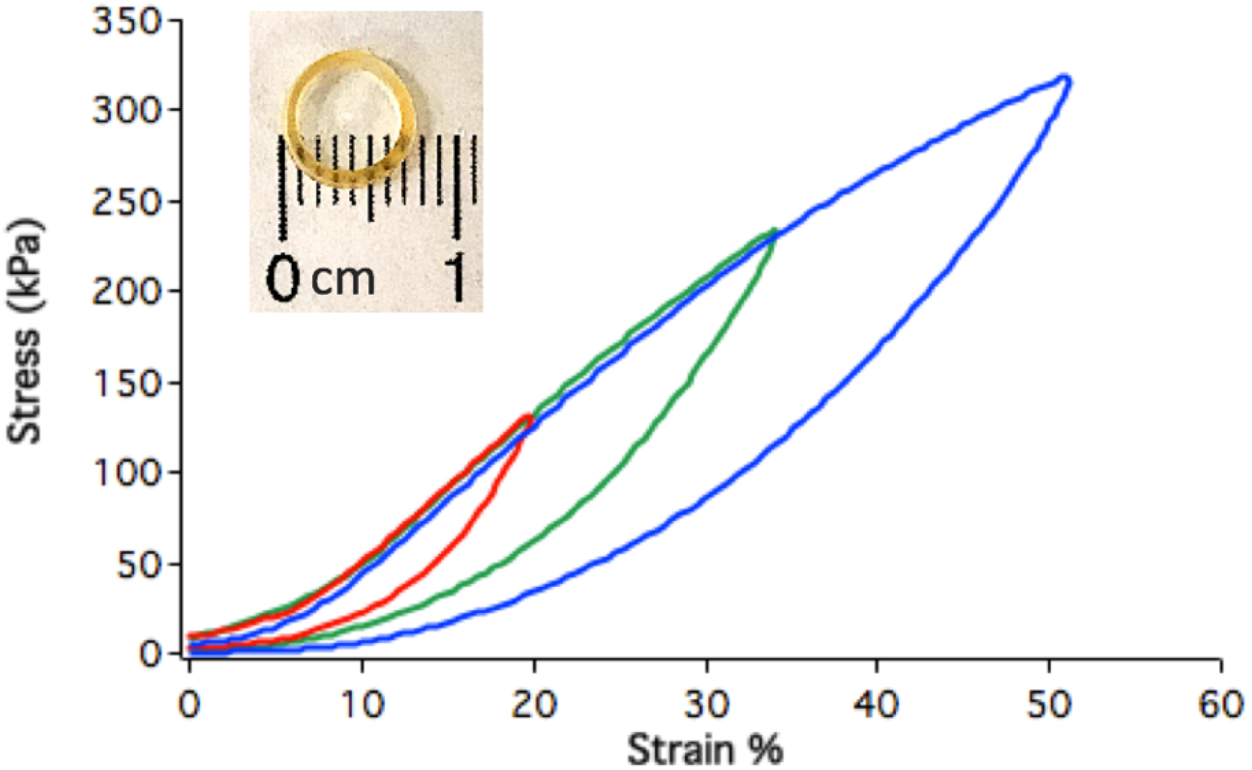
Typical tensile stress-strain curves of 20% N-DC (FM23C)_8_ hydrogels. The mechanical properties of (FM23C)_8_ hydrogels are similar to those of (FL)_8_ hydrogels. The Young’s modulus is 0.89 MPa. Inset shows an optical photograph of the N-DC (FM23C)_8_ hydrogel.

**Figure S7.**
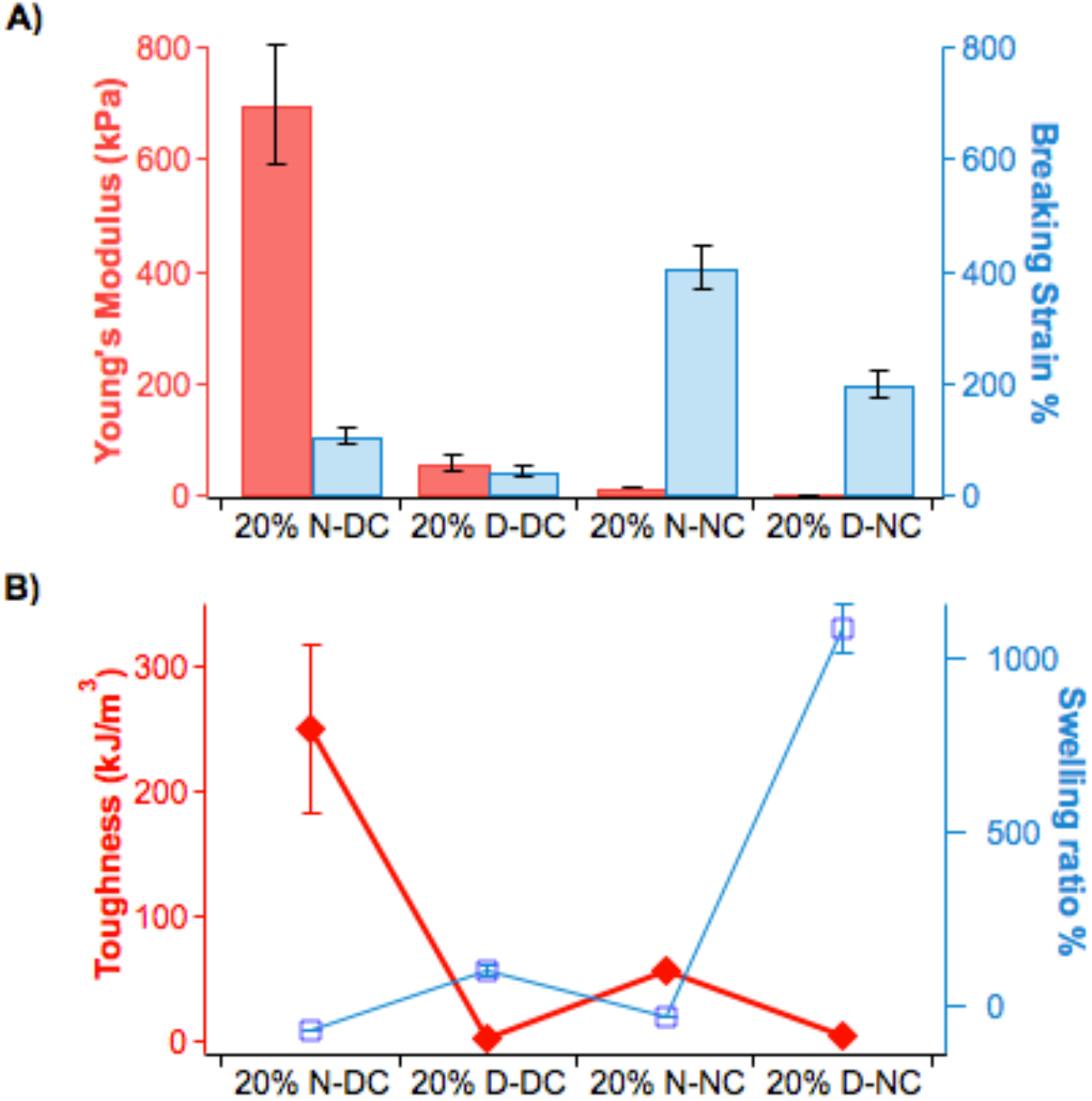
Mechanical properties of N-DC and D-DC (FL)_8_ hydrogels in tensile testing. A) Young’s modulus and breaking strain of N-DC and D-DC (FL)_8_ hydrogels. B) Toughness and swelling ration of N-DC and D-DC (FL)_8_ hydrogels. It is evident that the N-DC hydrogel exhibited much higher Young’s modulus and higher toughness than the N-NC hydrogel.

**Figure S8.**
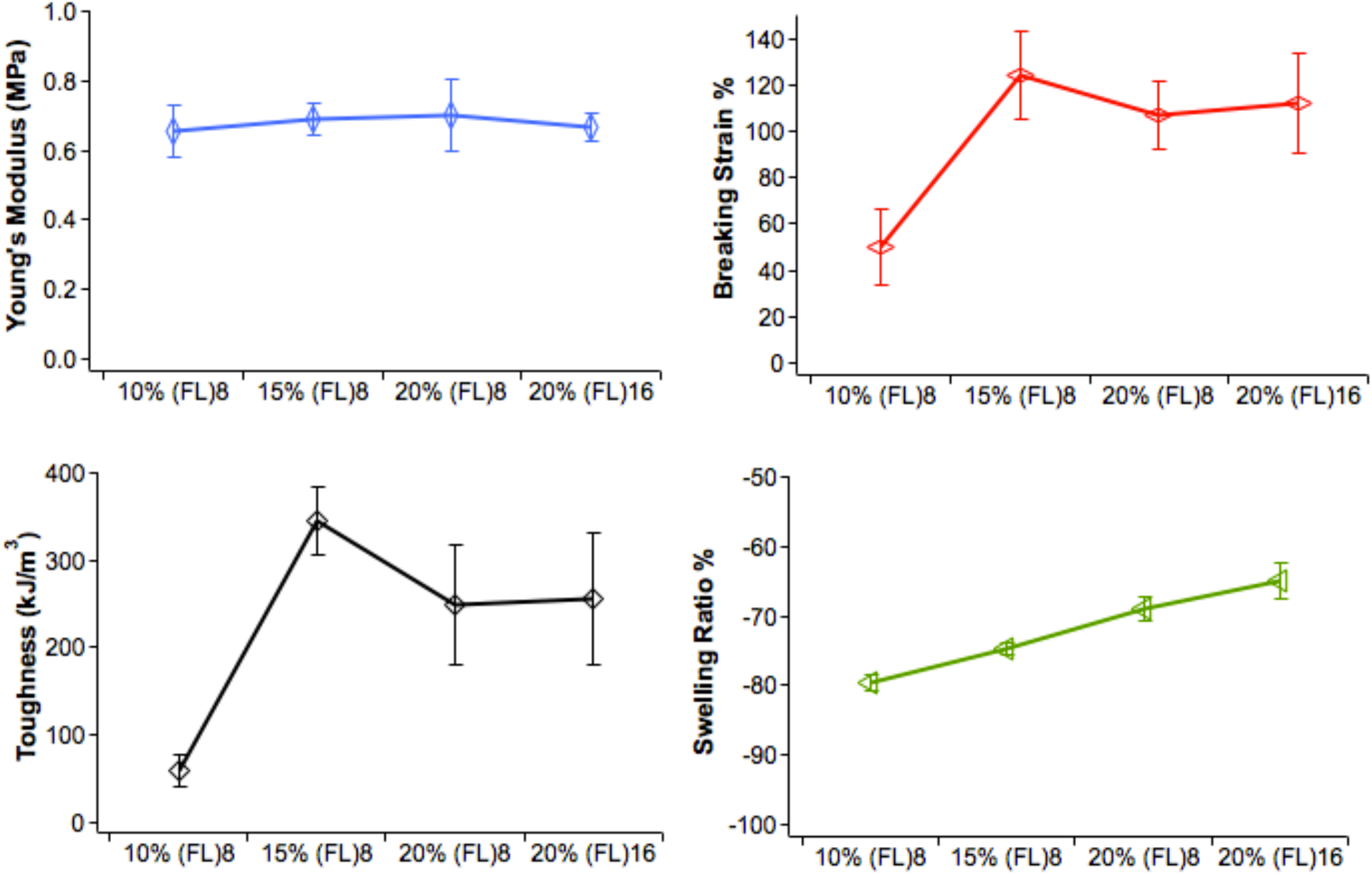
Tensile properties of N-DC (FL)_8_ hydrogels at different protein concentrations.

**Figure S9.**
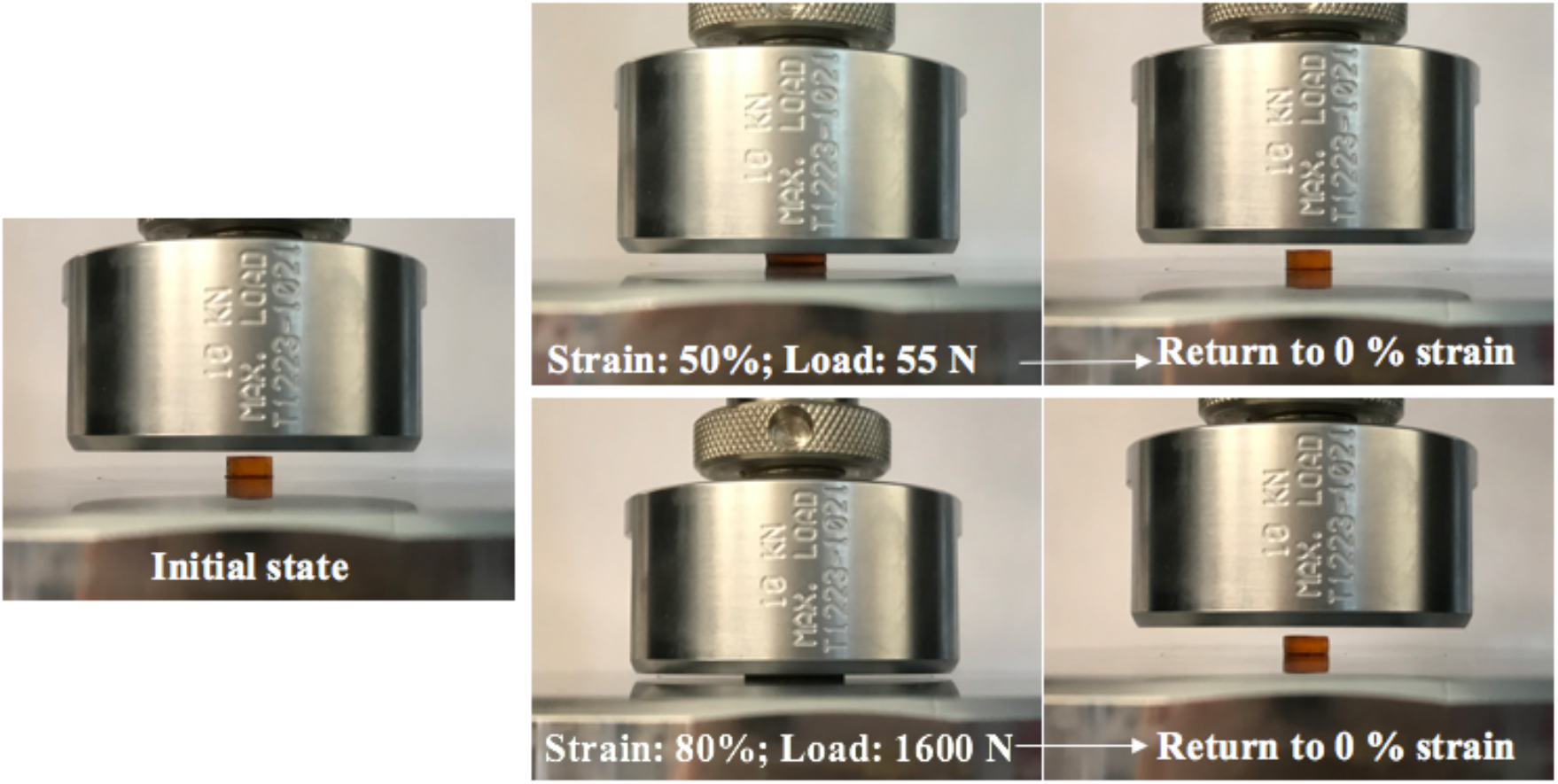
Photographs of the N-DC (FL)_8_ hydrogel under compression. After unloading, the hydrogel recovered its shape rapidly.

**Figure S10.**
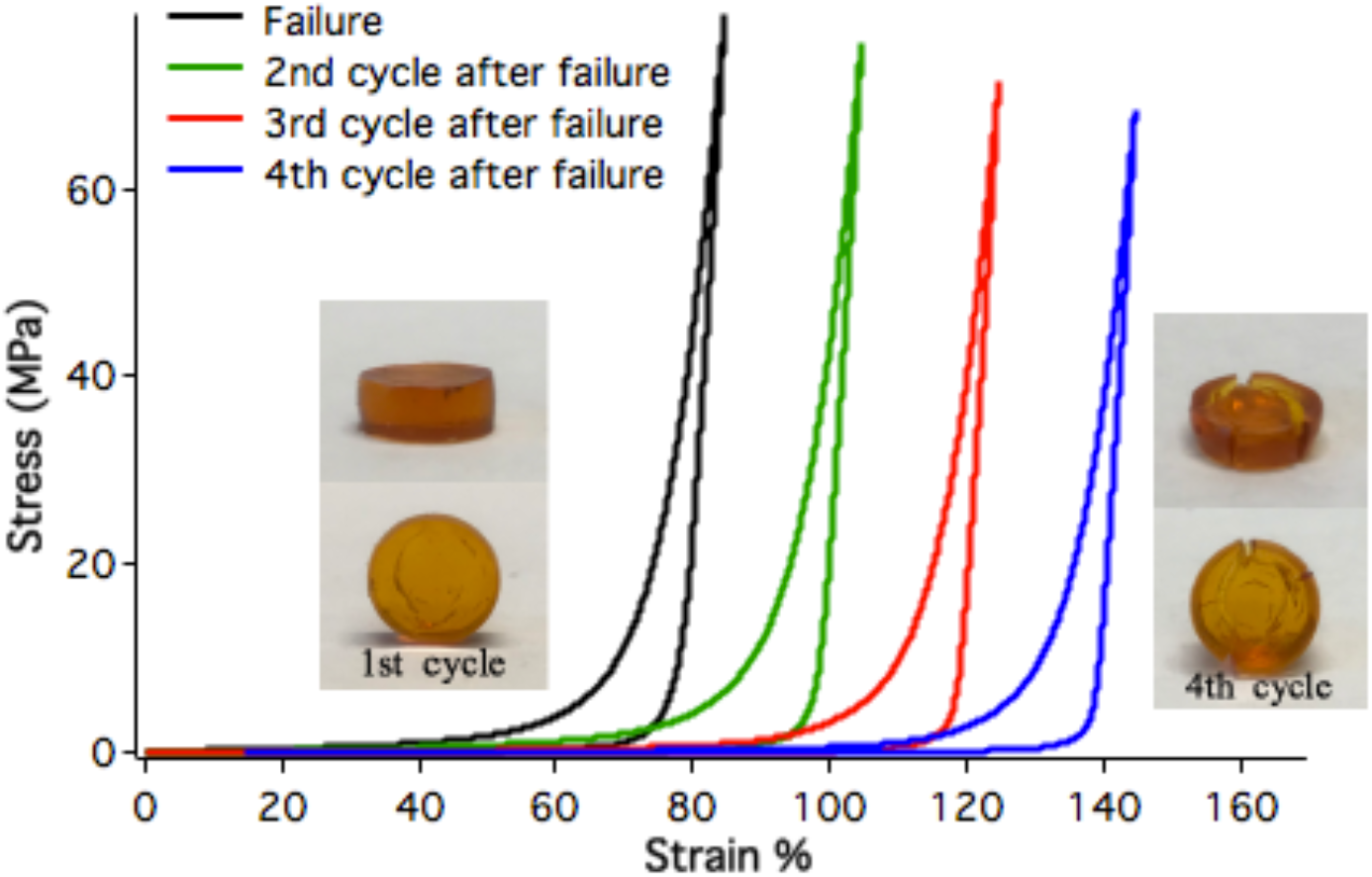
Stress-strain curves of a N-DC (FL)_8_ hydrogel compressed to failure. Inset shows the photographs of the hydrogel right after failure (1^st^ cycle) and after three more consecutive compression-unloading cycles (4^th^ cycle). Cracks were observed right after the failure. Subsequent compression led to the propagation of the crack.

**Figure S11.**
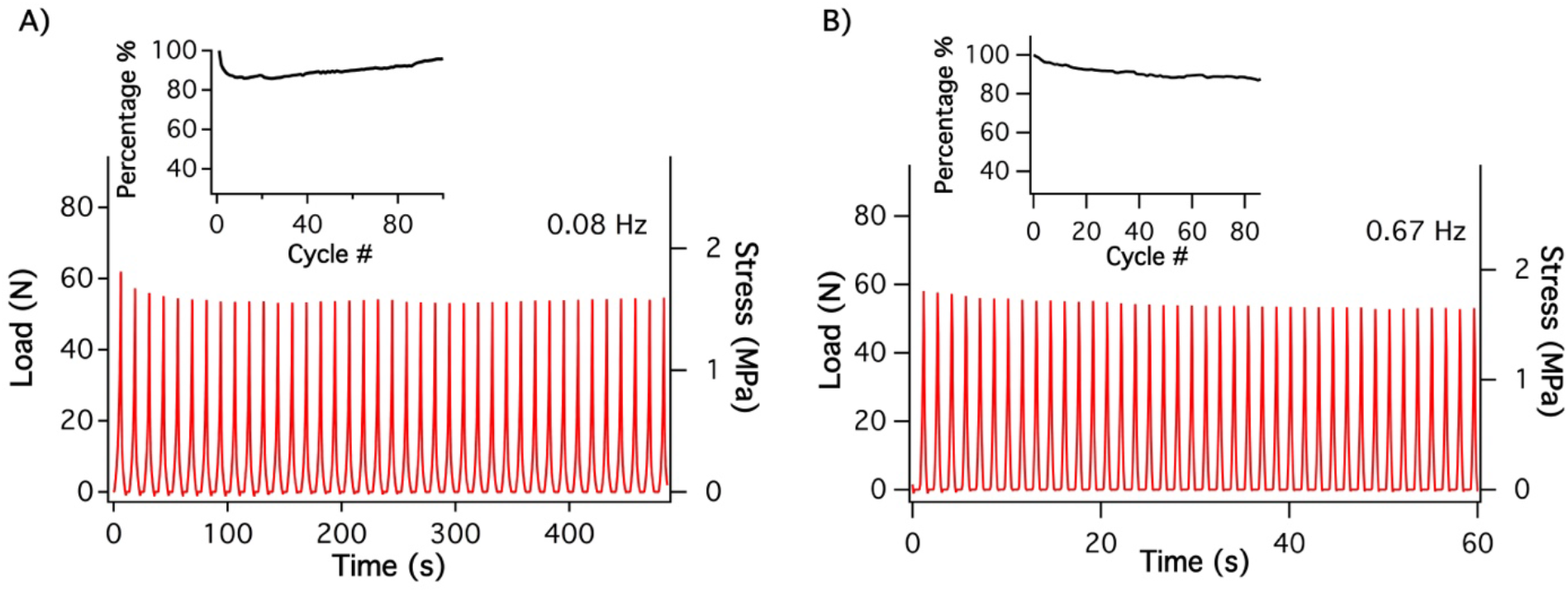
Consecutive compression-unloading curves of a N-DC (FL)_8_ hydrogel at a frequency of 0.08 Hz (A) and 0.67 Hz (B). The loading rate was 20 mm/min in A) and 200 mm/min in B), respectively.

Supplementary Movie 1. Consecutive compression-unloading cycles of the N-DC (FL)_8_ hydrogel. The final strain was 65%.

**Table S1.**
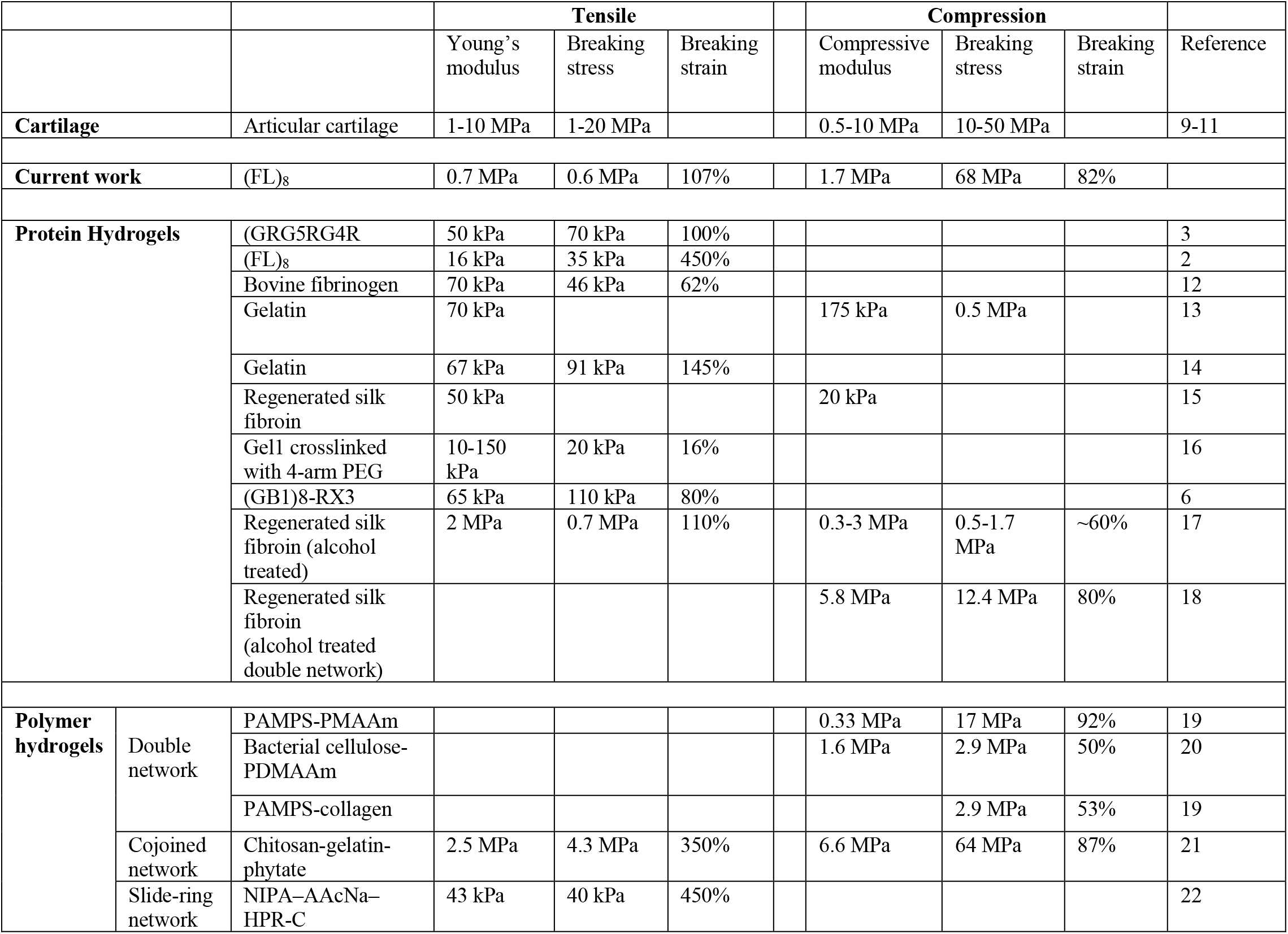
Mechanical properties of the state-of-the-art protein hydrogels and polymer hydrogels.

**Table S2.** Enhancement of mechanical properties via the DC hydrogelation method. (*E*: tensile modulus; *Y*: compressive modulus)

|  | Tensile |  |  | Compression |  |  |
| --- | --- | --- | --- | --- | --- | --- |
|  | N-DC hydrogel | N-NC hydrogel |  | N-DC hydrogel | N-NC hydrogel |  |
| Protein | $E_{N-DC}$ (kPa) | $E_{N-NC}$ (kPa) | Enhancement $E_{N-DC}/E_{N-NC}$ | $Y_{N-DC}$ (kPa) | $Y_{N-NC}$ (kPa) | Enhancement $Y_{N-DC}/Y_{N-NC}$ |
| (FL)<sub>8</sub> | 700 ± 110 | 15.7 ± 1.8 | 44 | 1614 ± 201 | 40.1 ± 2.7 | 40 |
| (NuG2)<sub>8</sub> | 620 ± 93 | 11.2 ± 3.3 | 56 | 1086 ± 31 | 55.0 ± 11 | 20 |
| (GB1)<sub>8</sub> | 227 ± 30 | 4.8 ± 1.1 | 47 | 339 ± 36 | 68.3 ± 10.5 | 5.0 |
| GRG<sub>5</sub>RG<sub>4</sub>R | 123 ± 53 | 32.7 ± 4.2 | 3.8 | 220 ± 31 | 19.6 ± 3.9 | 11 |
| N<sub>4</sub>RN<sub>4</sub>RNR | 126 ± 21 | 22.1 ± 2.7 | 5.7 | 592 ± 148 | 114.0 ± 25.1 | 5.2 |
| BSA | 236 ± 5 | 6.5 ± 0.1 | 36 | 776 ± 173 | 19.5 ± 3.9 | 40 |
| (GA)<sub>8</sub> | 438 ± 55 | 60.1 ± 9.0 | 7.3 | 1704 ± 248 | 95.0 ± 17 | 18 |

(NuG2)<sub>8</sub>: a polyprotein made of eight tandem repeats of the protein NuG2 (ref. 23)
(GB1)<sub>8</sub>: a polyprotein made of eight tandem repeats of the protein GB1 (ref. 24)
GRG<sub>5</sub>RG<sub>4</sub>R: G represents GB1 domain, R represents the 15 residue consensus sequence of resilin (ref. 3)
NRN<sub>4</sub>RN<sub>4</sub>R: N represent NuG2 domain, R represents the 15 residue consensus sequence of resilin
BSA: bovine serum albumin (ref. 25).
(GA)<sub>8</sub>: a polyprotein made of eight tandem repeats of the protein GA (ref. 26)

